# *SRGAP2* and the gradual evolution of the modern human language faculty

**DOI:** 10.1101/143248

**Authors:** Pedro Tiago Martins, Maties Marí, Cedric Boeckx

## Abstract

In this paper we examine a new source of evidence that draws on data from archaic human genomes to support the hypothesis that vocal learning in *Homo* preceded the emergence of Anatomically Modern Humans. We build our claim on the evolutionary history of the SLITROBO GTPase 2 gene (*SRGAP2*). The SLIT-ROBO molecular pathway has been shown to have an important role in the context of vocal learning. Though the relevance of the *SRGAP2* gene duplication in the emergence of some aspect of language has not gone completely unnoticed, recent results now allow us to articulate a mechanistic hypothesis of its role in the context of axon guidance. Specifically, *SRGAP2C*, a duplication of *SRGAP2* crucially also found in Neanderthals and Denisovans, but not in extant mammals, inhibits the ancestral *SRGAP2A*, which in turn modulates the axon guidance function of the SLIT-ROBO molecular pathway. This, we claim, could have contributed to the establishment of the critical cortico-laryngeal connection of the vocal learning circuit. Our conclusions support the idea that complex vocal learning could already have been part of the arsenal of some of our extinct ancestors.

## 1 INTRODUCTION

There has been much controversy among scholars regarding when the faculty of language arose in the evolutionary history of our species. Proposals put forward in the last decades cover a range of dates as large as 100,000-500,000 years ago [1, 2, 3, 4]. A recent special issue on the biology and evolution of language also reflects the disparity of competing positions [5].

Part of the problem when addressing this question lies in the fact that many researchers continue to see the language faculty as a homogeneous organic object. But we believe that it is far more promising, from a biological point of view, to see our linguistic competence as a complex mosaic formed by a species-specific (‘novel’) combination of several inherited and phylogenetically heterogeneous traits, tinkered with along traditional Darwinian lines [6, 7]. We expect many of these pieces of the language mosaic to be fairly straightforwardly recognized in other species (homologies), whereas other pieces may have less transparent roots [8]. Inasmuch as the appearance and development of these various traits is directly related to genetic factors, a crucial source of evidence for tracing the phylogenetic history of language, and ultimately timing its emergence, comes from the study of the genetic material remaining in fossils of ancient organisms. Progress in paleogenetics has dramatically changed the testability of some evolutionary scenarios [9]. A famous example of this was given in 2007 by Krause et al. [10], who found that *FOXP2*, a gene associated with language impairments and hampered orofacial movements [11], has the same two unique mutations in both Neanderthals and humans, critically missing in our closest extant great ape relatives. To the extent that these two mutations contributed to the establishment of some aspects of our brain’s language-readiness [12, 13], Krause et al.’s discovery strongly suggests that aspects of our language faculty had evolved prior to the divergence of the two lineages, some 600,000 years ago [14]. In this paper we focus on the evolutionary history of *SRGAP2*, which codes for the SLIT-ROBO Rho GTPase activating protein 2 (SRGAP2). We offer, on the basis of what we have learned from other species about vocal learning, another argument in support of the idea that vocal learning was established in *Homo* before the emergence of anatomically modern humans. While the link between *SRGAP2* duplication and language evolution has been mooted before [15, 16], we show how it has now become possible to provide a mechanistic articulation of this link, making the hypothesis fully testable.

### 1.1 Vocal learning in birds: a mirror for human language evolution

Vocal learning is the ability to learn to reproduce communicative signals from conspecifics. Such an ability is displayed in a limited number of lineages phylogenetically scattered across some groups of mammals (bats, elephants, cetaceans, pinnipeds, and humans) and birds (songbirds, parrots, and hummingbirds) [17, 18]. Among the pieces interlocked in the language mosaic, we have decided to focus on vocal learning here because it is the best understood to date in light of the recent literature [19, 20, 15]. As such, it provides the best testing grounds for evolutionary scenarios concerning some important aspects of human language.

The vocal learning literature, especially the line of research pursued by Erich Jarvis and colleagues, already offers interesting scenarios to test. Let us briefly sketch them here, as they will play an important role in the background of the next sections. Vocal learning birds and humans share a number of forebrain structures specialized in song and speech control, respectively [20]. Among them, all three learning avian species exhibit several brain nuclei that are distributed in two pathways: the anterior, or vocal learning pathway, which is mainly specialized in vocal imitation and malleability, and the posterior, or vocal production pathway, which associates with the intentional production of (learned) vocalizations. Within this posterior pathway, which will be the main focus in the following sections, oscines, parrots, and hummingbirds present three analogous motor regions in the cortex, namely the robust nucleus of the arcopallium (RA), the central nucleus of the anterior arcopallium (AAC), and the vocal nucleus of the arcopallium (VA), respectively, which are in turn analogous to the laryngeal motor cortex (LMC) in humans. In both learning birds and humans, this nucleus makes a direct projection to the brainstem motor neurons (MN) that control the syrinx in birds and the larynx in humans [21, 15, 22, 20, 23].

On the basis of such similarities, a motor theory of vocal learning has been proposed [22], arguing that cerebral systems specialized for vocal learning in distantly related lineages are independent evolutions of a motor system inherited from their common ancestor. Analyses in gene expression [22, 24, 25, 26] certainly point in this direction, further supporting that the posterior pathway, which we will focus on next, must have emerged from a primitive motor system [15, 22, 27]. Since several forebrain motor learning pathways with sensory input appear to be formed during early development by successive duplications, thereafter projecting to various brainstem or spinal cord neurons associated with different muscle groups, it has been proposed that the posterior connection appeared similarly as one further duplication that then projected to the brainstem MN in charge of the vocal organs [28, 15]. Pathway duplication unfolds in a manner analogous to gene duplication—with a whole pathway duplicating and the duplicate taking on new function—, and actually having gene duplication as one possible underlying mechanism [29].

Neuroanatomical research conducted with primates has identified homologous representations of the larynx in the motor cortex (LMC) both in human [30, 31] and in non-human primates, such as chimpanzees (*Pan troglodytes*) [32], rhesus monkeys (*Macaca mulatta*) [33, 34], and squirrel monkeys (*Saimiri Sciureus*) [35, 33]. However, although the LMC connectivity network is broadly similar among primates tested, a robust cortico-laryngeal direct projection to the vocal MN in the brainstem has been found only in humans [23, 36].

There are reasons to believe that the posterior pathway develops gradually, as it is present at a very rudimentary level in the brain of a non-vocal-learning suboscine species. Indeed, as Liu et al. have shown [37], the eastern phoebe (*Sayornis phoebe*), closely related to songbirds, possesses a specialized forebrain region that seems homologous to the RA in oscines. This region presents descending projections to the brainstem respiratory nucleus and has a singing-associated function. In this regard, eastern phoebes present a long period (8-9 months) of song plasticity before its crystallization. This circuitry seems to be a proto-form of what we find in vocal learning oscines, though not developed enough for vocal learning brain-readiness inasmuch as, unlike in songbirds, there is no direct projection from the arcopallial RA-like nucleus to the tracheosyringeal neurons.

Once this critical neural pathway is established, it is quite likely to undergo further elaborations, giving rise to more complex forms of vocal learning. A case in point that can serve as an example for such specializations can be found in parrots, known to be able to imitate vocalizations of conspecifics, but also sounds produced by other species. A study involving the three superfamilies of parrots (*Strigopoidea*, *Cacatuoidea*, and *Psittacoidea*) [38] has revealed an internal subdivision in their song cortical nuclei, wherein a core region shows different gene expression from the surrounding shell area, while both exhibit in turn different expression from the surrounding motor cortical region. Interestingly, the posterior connection to the brainstem MN associated with the syrinx, along with other connections with different forebrain vocal regions, is projected exclusively from the core region and not from the shell [15]. Chakraborty and Jarvis [29] suggest that the core region in the parrot AAC evolved convergently in all three avian vocal learning species via duplication from the surrounding motor regions, and subsequently the shell area was developed in parrots, allowing for their more complex vocal proficiency.

As we just saw, critical neural stuctures such as the posterior pathway, taken as a reference point for the origin of the vocal learning capacity, likely emerge in proto-form, and, once present, can be subject to further elaboration, under the influence of several factors. We believe that the same could be true for the emergence of language in our lineage [39].

### 1.2 The *SRGAP2* gene suite and the timing of critical evolutionary steps in Homo

Although *SRGAP2* is highly conserved among mammals [40] and has remained unchanged at least in the last 6 million years of our evolution (its F-BARx domain is identical in humans, chimpanzees, bonobos, and orangutans) [41], it has given rise to three human-specific duplications, two of which underwent subsequent mutations. The sequence of events, identified by Dennis et al. [40], illustrated in Fig. 1, happened as follows (the chronological ranges have been calculated assuming the timing of divergence between chimpanzee and human lineages within a span of 5-7 million years ago (mya), based on fossil records [42, 43, 44] and genetic analyses [45]): The first duplication took place 2.8–3.9 mya, when the promoter and first nine exons of the original gene—designated *SRGAP2A* to distinguish it from its derivatives— were duplicated from the locus 1q32.1 to 1q21.1, thus giving rise to the primitive *SRGAP2B* (*P-SRGAP2B*). A second duplication occurred 2.0–2.8 mya, when *P-SRGAP2B* was copied from 1q21.1 to 1p12, leading to the primitive *SRGAP2C* (*P-SRGAP2C*). In the aftermath of this event [40, 41], the two primitive duplicated copies, *P-SRGAP2B* and *P-SRGAP2C*, accumulated non-synonymous mutations which resulted in the contemporary *SRGAP2B* and *SRGAP2C* forms, carrying five (R73H, R108W, R205C, R235H, R250Q) and two (R79C, V366L) aminoacid replacements, respectively. Finally, the third and last duplication, which occurred 0.4-1.3 mya, copied the modern *SRGAP2B* within 1q21.1, thus giving rise to *SRGAP2D* [40]. Consistent with the timing of their appearances, all three human paralogs, *SRGAP2B*, *SRGAP2C*, and *SRGAP2D*, have been found also in the genomes of Neanderthals and Denisovans [46].

**Figure 1:**
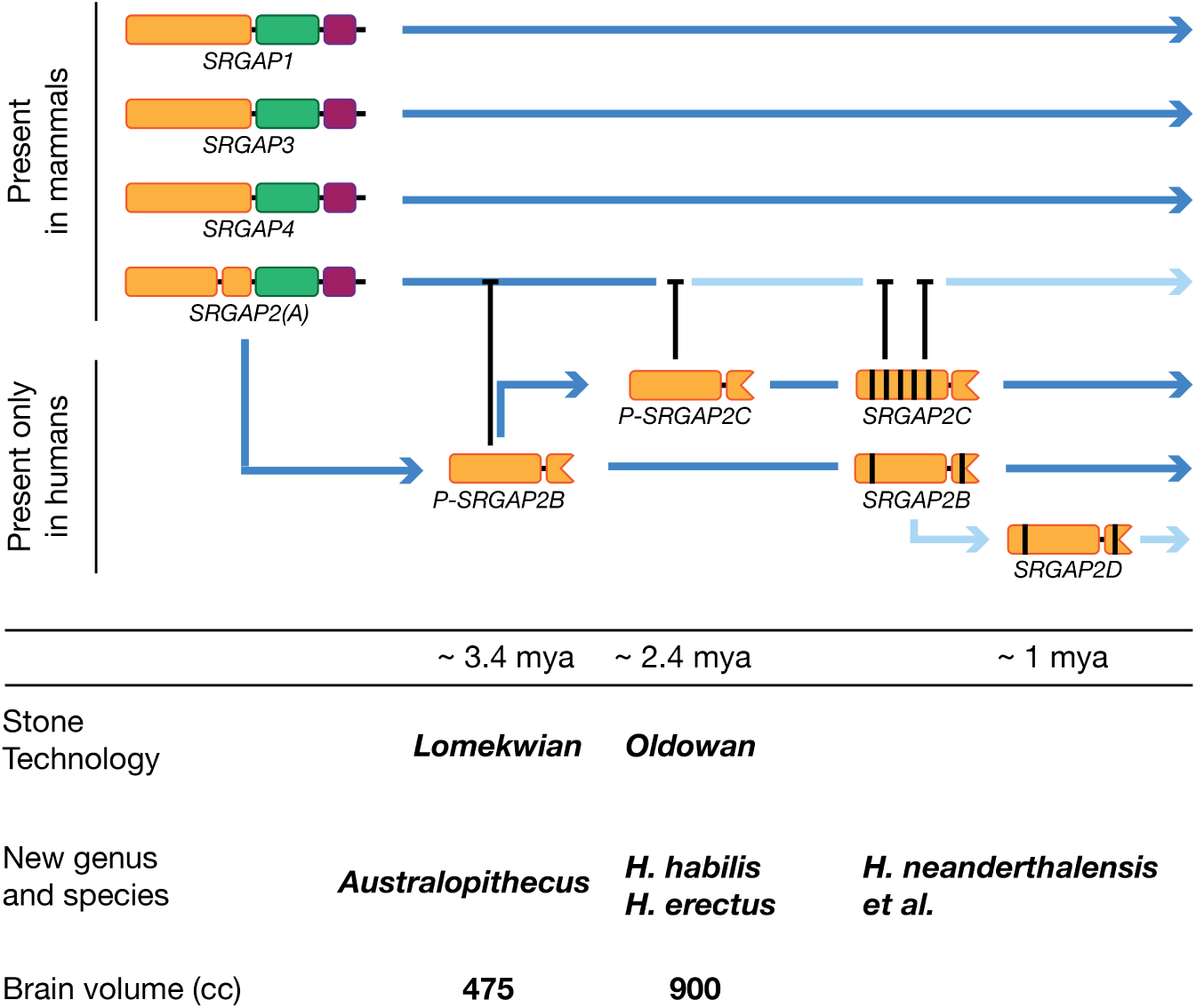
Evolutionary history of SRGAPs and chronological coincidences with human landmarks. On top, the colored figures represent each of the SRGAP genes. In orange, the F-BAR domains, with an F-BAR extension in the case of *SRGAP2(A)*. The human duplicate copies are devoid of RhoGAP (green) and SH3 (violet) domains, but conserve the most part of the F-BARx domain. Blue arrows symbolize functional continuity of the gene; the reduced activity of *SRGAP2(A)* by *SRGAP2C*, along reduced activity of *SRGAP2D*, are represented by arrows in a lighter shade of blue. The dates in the central horizontal fringe correspond to the emergence of the primitive form of *SRGAP2B* (*P-SRGAP2B*; ~ 3.4 mya) and *SRGAP2C* (*P-SRGAP2C*; ~2.4 mya), which parallels the first (Lomekwian) and second (Oldowan) known generations of stone technology. The aminoacid replacements that *P-SRGAP2B* and *P-SRGAP2C* underwent to reach their modern forms (two in *SRGAP2B*, five in *SRGAP2C*) are represented by black bars. Around ~1 mya, *SRGAP2D* emerged as a copy of *SRGAP2B* and carries the same two substitutions. The penultimate row in the figure gives account of the chronological coincidences between the duplication events that led to *P-SRGAP2B* and *P-SRGAP2C*, and the appearances of the genus *Australopithecus* and *Homo* (*H. habilis*; *H. erectus*), respectively; similarly, the appearance of *H. neanderthalensis*, likewise that of other sister *Homo* species, parallels the emergence of *SRGAP2D*. At the last row, the differences between the estimated brain size of *Australopithecus* (475 cc) and those of *Homo habilis* and *Homo erectus* (900 cc).

Importantly, the timing of the *SRGAP2* duplications appears to correspond fairly closely to some landmarks in our lineage in terms of brain size and use of stone tools in the transition from *Australopithecus* to *Homo*, raising the possibility that the relevant duplications contributed to these phenotypic changes [47, 16, 39]. Thus, the time of the first duplication (*P-SRGAP2B*) matches the appearance of *Australopithecus*, which had an average brain size of ca. 475 cc, similar to that of genus *Pan*. The second duplication span (*P-SRGAPC*) corresponds to the appearance of *Homo habilis* and *Homo erectus*, having an average brain size of ca. 900 cc. Finally, the last duplication (*SRGAP2D*) is associated with the emergence of late *Homo erectus*, of Neanderthals and of other sister species [46]. In addition, the timing of the first and the second duplications, *P-SRGAP2B* (*~*3.4 mya) and *P-SRGAP2C* (*~*2.4 mya), shows a fairly close correspondence with the first and second generations of the use of stone tool technology, Lomekwian and Oldowan [41].

In light of claims that total number of neocortical neurons is shown to be a better correlate of cognitive complexity than brain size per se (both absolute or relative) [48, 49], it is also interesting to point out that the evolutionary rate of the *SRGAP2* gene has been claimed to positively correlate with an increase in the number of cortical neurons in mammals [50].

Not surprisingly, several authors suggested that *SRGAP2* duplications may underlie some of the changes that led to human cognition. The most explicit suggestion along these lines that we are aware of was made in [15]. Building on the existing literature on the functional effects of the relevant duplications, Chakraborty and Jarvis [29] write:

> The duplicated copies act as competitive inhibitors to slow cortical dendritic development of already existing brain pathways, which in turn allow greater neural plasticity into adulthood. *SRGAP2* modulates activity of the ROBO axon guidance receptors, which are in turn activated by the SLIT family of protein ligands to modulate axonal/dendritic migration and branching in various brain regions. Intriguingly, the *SLIT1* ligand is uniquely downregulated in the song production nucleus RA analogue of vocal learning birds (songbird RA, parrot AAC and hummingbird VA) and the analogous human LMC, which would mean that there could be a synergistic effect of the duplicated *SRGAP2* GTPase and lower *SLIT1* levels in the duplicated vocal motor pathways in humans.” [references omitted]

We find this suggestion very insightful, and what follows is meant to provide support for it. Doing so requires spelling out some of the assumptions and findings that are alluded to in this quote. We turn to this next.

## 2 *SRGAP2* genes, filopodia, and axon guidance

The first thing to point out in the context of Chakraborty and Jarvis’ suggestion is that the existing literature on *SRGAP2* does not immediately support it. Despite their names (*SRGAP* genes —*SLIT-ROBO GTPase activating protein* coding genes), the nature of the interactions between *SLIT* genes, *ROBO* genes, and *SRGAP* genes does not always go in the desired direction for vocal learning, by which we mean the axon guidance role, for reasons we discuss briefly in the next subsection.

### 2.1 *SLIT* and *ROBO* axon guidance genes and the vocal learning posterior pathway

As it has been said above, a direct neural projection from a cortical/pallial motor nucleus and the brainstem MN controlling the larynx/syrinx appears to be a key component in the evolution of the vocal learning ability. To form this structure during the early development of the brain, the axonal extensions of the neurons in the cortical region must be sent and guided along pathways to eventually reach their synaptic targets in the brainstem through a process which requires the action of axon guidance genes [51].

In this regard, as alluded to in the quote from [15], studies conducted with birds from the three groups of species of avian vocal learners [52, 21], have shown that axon guidance genes of the SLIT-ROBO families present a convergent differential regulation in the pallial motor nucleus of the learning species.

Summarizing briefly these results, we can say that *SLIT1*, a gene belonging to the SLIT family of repulsive axon guidance genes [51], shows a differential downregulation precisely in the songbird RA and in the analog regions in parrots (AAC) and hummingbirds (VA), i.e. the arcopallial nuclei making the direct projection to the brainstem MN. The expression of *SLIT1* in these nuclei is remarkably low compared to the surrounding arcopallium. More precisely, in the case of the parrot AAC, which has a subdivision between core and shell we had already expounded, the downregulation of *SLIT1* occurs only in the core region, which is the one sending the projection to the brainstem MN. In contrast, no such regulation of *SLIT1* was observed either in the arcopallium of non-vocal learning birds tested (quails and ring doves) or in a recently discovered putative LMC of mice, thus highlighting the specificity of this expression pattern to vocal learning lineages [52]. All in all, the particular pattern of expression of *SLIT1* strongly suggests a functional relation between the downregulation of the axon guidance factor and the formation of the neural projection from the cortical nucleus to the brainstem MN in charge for the syrinx, a relation which would be consistent with the similar downregulation of *SLIT1* that has been found in the human LMC [21].

*ROBO1* belongs to the Roundabout (ROBO) family of axon guidance genes, whose encoded proteins act as receptors of SLIT ligands to transduce the repulsive cue into the intracellular domain [53, 51, 54]. As *SLIT1*, *ROBO1* also shows a differential expression in relation to the posterior pathway: upregulated in the parrot AAC core and in the hummingbird VA, compared to the surrounding arcopallium, whereas in the songbird RA it is downregulated. Despite the divergence in songbirds with respect to the other two groups, *ROBO1* has been observed to be temporarily upregulated in male zebra finches (endowed with a higher capacity for song compared to females) between posthatch days 35 and 65, a period deemed critical for vocal learning [52].

### 2.2 SRGAPs, SLITs, and ROBOs

In mammals, the SRGAP family of genes consists of four members: *SRGAP1*, *SRGAP2*, *SRGAP3*, and the distantly related *SRGAP4* [55]. The first three were uncovered in 2001 by Wong et al. [56] in a yeast two-hybrid experiment in which the SRGAPs were found to interact with the C-terminal region of rat *ROBO1*. After their identification, the researchers further analyzed, through different in vitro experiments in human embryonic kidney (HEK) cells, various aspects of the interaction between *SRGAP1* and *ROBO1*, including the effect of extracellular *SLIT2* in such binding. Among other results, they found that extracellular *SLIT2* upregulated *ROBO1* -*SRGAP1* binding in a dose-dependent manner, thus leading to the inactivation of *CDC42*, a member of the Rho GTPase family, which has a well-documented role in the regulation of the cytoskeletal dynamics [57]. In the light of these findings, the authors proposed that the newly discovered SRGAPs are intracellular effectors in the downstream of a SLIT-ROBO signaling pathway and play a role in the guidance function of SLITs. This approach would make it possible, therefore, for *SRGAP2* to interact with *ROBO1* downstream of an axon guidance cue, are part of the mechanism leading to the constitution of the aforementioned posterior pathway.

However, and disappointingly for our purposes, subsequent research did not provide support for this initial proposal concerning ROBO1-SRGAP2 binding. Building on the suggestion in [56], [58] investigated the SRGAPs mRNA expression in rat brain, at various developmental stages and could find only a relative coincidence with the localized *ROBO1* expression reported by other scholars [59, 60]. A subsequent study [61] on SRGAPs expression in several embryonic and postnatal stages noted similarities of SRGAP2 pattern with that of *ROBO2*, but did not report any interaction with *ROBO1*. Li et al. focused on the CC3 motif of *ROBO1* that [56] had found in interaction with the SH3 domain of *SRGAP1*, and then assessed their binding with the SH3 domains of *SRGAP1*, *SRGAP2*, and *SRGAP3* [62]. The result was that most of the recreated peptides did not bind, and only one showed a feeble and transient interaction. Similarly, [63] did not identify *ROBO1* as a ligand for *SRGAP2*. (Below we return to these unsuccessful attempts, as a recent study [64] provides a possible reason for these results.)

On a more positive note, SRGAPs, and specifically *SRGAP2* on which we focus here, have been reported to serve various functions regarding cortical development at early stages. First, *SRGAP2* has been shown to regulate axon-dendrite morphogenesis and neuronal migration through its ability to induce protrusions at the plasma membrane. A study of cortical neurons in mice showed that the knockdown of *SRGAP2* significantly decreased both dendritic and axonal branching, while, on the other hand, neurons with shRNA-silenced expression of *SRGAP2* migrated roughly 25 per cent faster than the control group, thus showing an inhibitory effect [65]. These results support the suggestion in [56] (based on experiments on *SRGAP1*) that SRGAPs can regulate cell migration. A subsequent study [66] showed the same effects in vivo, and demonstrated, in addition, that the expression of *SRGAP2C* in mouse cortical neurons had a similar effect to that caused bi-ancestral *SRGAP2* knockdown, viz. an increase in the rate of cell migration. In the knockdown condition, [66] added another function of *SRGAP2* to those already established: it promotes the maturation of the dendritic spines and limits their density. Indeed, an experiment in vivo carried out with heterozygous *SRGAP2*-knockout mice revealed a substantially higher density of dendritic spines by comparison with the control group, with thinner and longer spines. [66] also found that the expression of SRGAP2C in mouse pyramidal neurons inhibited the function of *SRGAP2A* and extended the period of development of the spines (spinal “neoteny”), thus evoking an increase in their number per unit area and in their length. Interestingly, this last trait is considered characteristic of the human neocortex [67], and led to claims linking *SRGAP2* duplication with this particular property of the human neocortex.

As a final remark on the function of SRGAPs, we report their ability to co-regulate the ratio between excitatory and inhibitory synapses at their early development to reach the correct equilibrium at the mature stage. A recent in vivo study [68] in mouse cortical pyramidal neurons has shown that *SRGAP2A* increases the growth of inhibitory synapses and restricts their density. Curiously, in a way similar to the one mentioned earlier for dendritic spines, *SRGAP2C* antagonizes functions of *SRGAP2A* during synaptic development, prolonging their maturation period and increasing their final density.

As a result, *SRGAP2* duplication has not figured prominently in the literature on the evolution of vocal learning, since to the best of our knowledge neotenous spines are not (yet) considered a central property of vocal learners. Other more established neural traits associated with vocal learning appear not to be directly connected with the role of *SRGAP2*. Nevertheless, in the following sections we show how the well-documented function of *SRGAP2*, namely its ability to regulate protrusions at the plasma membrane of the neuron [65, 69, 70, 41], can be related to more canonical properties of vocal learning-ready brains, specifically axon guidance.

### 2.3 *SRGAP2* and axon guidance: an indirect link

In the process of axon guidance, a series of secreted proteins, such as the SLIT family, act as extracellular biochemical guiding effectors by evoking a signaling cascade that ultimately changes the cytoskeletal dynamics of the axon and directs its outgrowth either towards or away from the signaling source. These directional changes take place at the growth cone, a motile structure located at the distal end of the axon which is endowed with two types of F-actin–based structures: filopodia, which are narrow cylindrical protrusions based in unbranched parallel bundles of actin filaments (F-actin) formed by Ena/VASP and formin proteins, and lamellipodia, sheet-like protrusions based in a network of branched actin which is formed by the Arp2/3 complex. Axon guidance can be understood as a directed, recurrent process of enlargement and maturation of the growth cone, starting with the formation and extension of filopodia and lamellipodia at its leading edge, through the polymerization of actin filaments, followed by the flow of filopodia along the sides of the growth cone. The final step of the process is their eventual retraction at the base of the growth cone caused by the depolymerization of the F-actin. This last retraction allows the membrane to contract, thus forming a cylindrical consolidated axon shaft [71, 51]. Although the mechanisms whereby axons manage to find the correct pathways across the nervous system remain to be fully characterized, the two actin-supported structures that are characteristic of the axon growth cone, filopodia and lamellipodia, are considered to play a crucial role [71].

In relation to filopodia and axon guidance, a recent study in vivo in mouse dorsal root ganglia cells [72] has investigated the dynamics of the growth cone specifically during the axonal repulsion evoked through the SLIT-ROBO molecular pathway. Crucially for us, it has reached an unexpected conclusion: despite the classic view whereby a repulsive signal entails actin depolymerization at the side of the growth cone facing the guidance source, the amino-terminal fragment of *SLIT2* that contains the domain responsible for binding to *ROBO1* and *ROBO2* induced the formation and elongation of actin-based filopodia at the axon growth cone via SLIT-ROBO molecular pathway. Importantly, these SLIT-induced filopodia, which are longer and elongate distinctively toward the sources of the repulsive cue, are indispensable to elicit the guiding signal in the downstream of SLIT-ROBO. We think that these results are essential to understand how *SRGAP2A*, and perhaps some of its human-specific paralogs, can be related to axon guidance (see Fig. 2), thus supporting Chakraborty and Jarvis’ suggestion [29], and enabling us to provide novel support for the claim that vocal learning was established fairly early in our lineage.

**Figure 2:**
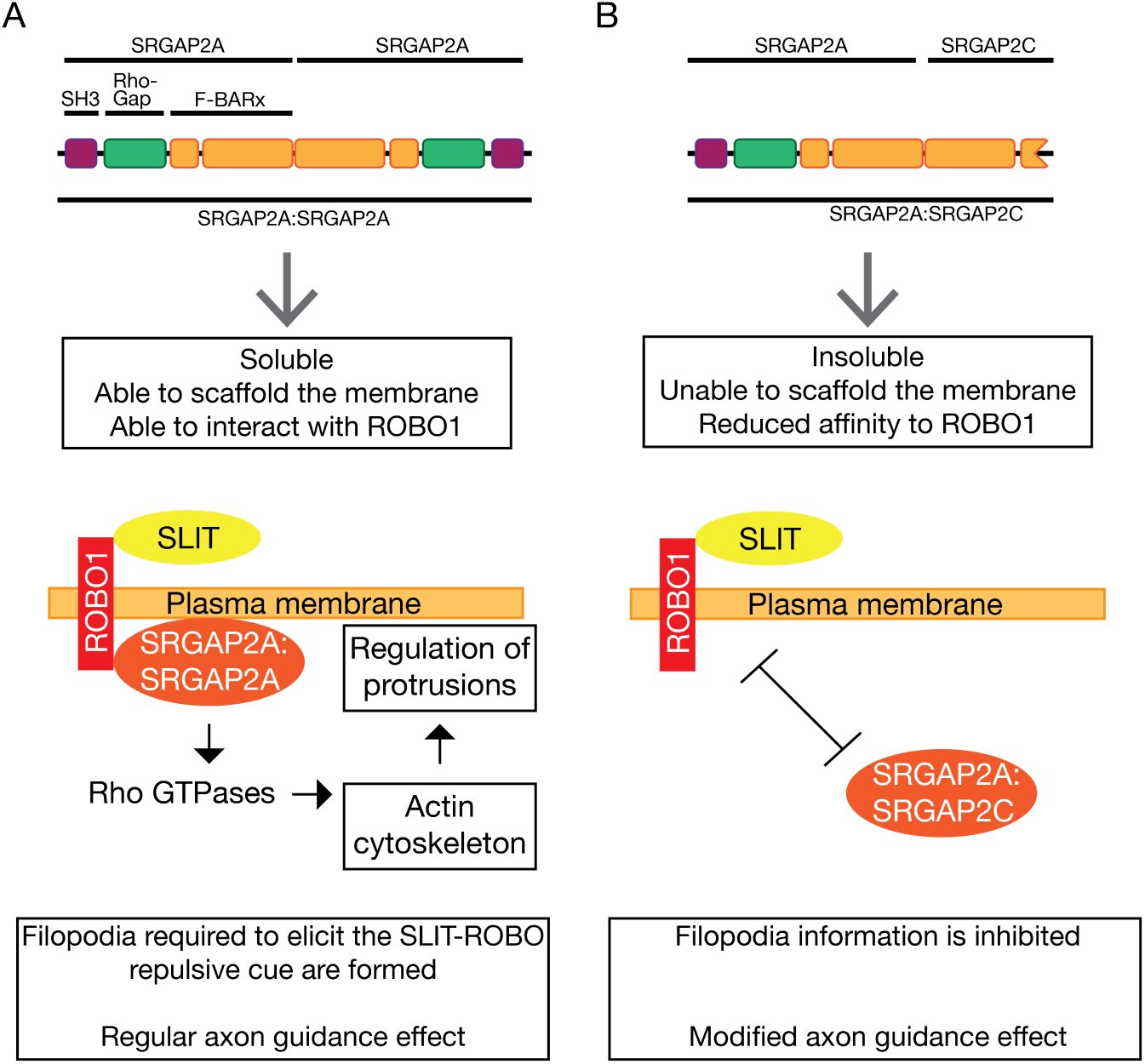
Proposed model for the implication of SRGAP2A and SRGAP2C in an axon guidance signaling pathway. A: SRGAP2A molecules homodimerize through their F-BARx domains, thus forming soluble dimers. These dimers have a singular inverse geometry which allows them to colocalize at the membrane at sites of protrusions. Once in place, these molecules are able to transduce a SLIT-ROBO axon guidance cue by interacting with Rho GTPases through their RhoGAP domains, thus regulating the actin cytoskeleton and scaffolding protrusions. The chain of interactions leads to the constitution of filopodia which extend towards the sources of SLIT. These filopodia are crucial to elicit the repulsive axon guidance cue. B: SRGAP2C heterodimerizes with SRGAP2A. The resulting molecule is insoluble, unable to scaffold the membrane, and has a limited affinity for ROBO1. Thus, SRGAP2C inactivates SRGAP2A’s ability to regulate filopodia, ultimately resulting in a modified effect in axon guidance.

### 2.4 SRGAP2A and SRGAP2C

*SRGAP2A* has a singular threefold composition: an F-BAR domain, which has an amino-terminal extension; a RhoGAP domain, and an SH3 domain [41]. Remarkably, the extended F-BARx domain allows the protein to explore the geometry of the membrane and to bind selectively to bulging sites or protrusions [69, 65, 70]. Once in place, *SRGAP2A* can regulate the dynamics of the actin-based cytoskeleton through its RhoGAP domain, thus evoking different effects in these protrusions. As examples of this, Guerrier et al. [65] showed that the overexpression of the *SRGAP2A* F-BAR in cortical neurons induced filopodia-like membrane protrusions, whereas Fritz et al. [70] have shown that it evoked a retraction of the membrane protrusions in a cell-cell overlap context by inactivating local pools of *Rac1* and *CDC42* which, in turn, caused a breakdown of the actin-supported cytoskeleton and the subsequent retraction. There may be several factors conditioning the specific result of the protrusion regulation that *SRGAP2A* evokes, but, as Fritz et al. note [70], one of them must be the upstream input that it receives, most likely from the SLIT-ROBO pathways. In fact, they show that the detected effect of *SRGAP2A* is elicited in the downstream of the *SLIT2* -*ROBO4* signaling. It is in the context of binding axon guidance molecules that the SH3 domain has shown to be indispensable, although not exclusive, since all three domains (F-BARx, Rho-GAP, and SH3) have been proven to exert a cooperative participation in binding *ROBO1* [64]. As Guez-Haddad et al. point out [64], this must be the reason why previous attempts to attest a significant interaction between *ROBO1* and the isolated SH3 domain of *SRGAP2A* (summarized above) had failed. Summing up then, the particular threefold composition of *SRGAP2A* endows it with the ability to regulate membrane protrusions likely in the downstream of the axon guidance SLIT-ROBO pathway.

SRGAP2A molecules are homodimers in solution. Prototypically, F-BAR domains form anti-parallel dimers that bind the plasma membrane through their concave N-surface, thus associating with membrane invaginations. However, the SRGAP2A homodimerization is not only mediated by the F-BAR domain, as typically could be expected, but rather by a large interface that includes the F-BAR, its Fx extensions, the RhoGAP, and the SH3 domains. This particular cooperative dimerization, which additionally increases the ability of the dimer to bind the membrane, evokes an inverted, convex N-surface that associates with protrusions instead of invaginations. The potential of *SRGAP2A* to regulate membrane protrusions likely depends on this particular form of homodimerization [41].

The duplicated copy *SRGAP2C* consists of a truncated form of *SRGAP2A* containing nearly all of the F-BARx with three modifications, two of which occurred in the first duplication event (around 3.4 mya), thus being present in the primitive forms, *P-SRGAP2B* and *P-SRGAP2C*. As Sporny et al. [41] have recently shown, SRGAP2C has the ability to heterodimerize with SRGAP2A, a property which was already present in the primal form *P-SRGAP2C*, which appeared *~* 2.4 mya. Crucially, unlike SRGAP2A homodimers, SRGAP2A:SRGAP2C heterodimers are insoluble, thus being unable to reach the proper sites in the plasma membrane and consequently being rendered inactive. An experimental quantification of the effect of *P-SRGAP2C* and *SRGAP2C* in compromising *SRGAP2A* solubility has been carried out by Sporny et al., reflecting that, when coexpressed with recreated P-SRGAP2C and with SRGAP2C in Sf9 cells, 60 and 40 per cent of *SRGAP2A* respectively were insoluble. In light of these data, it is clear that SRGAP2C acts as an inhibitor of SRGAP2A by cancelling its ability to bind to the membrane and regulate protrusions. Relevantly, this capacity of SRGAP2C to form stable heterodimers with SRGAP2A and its consequent efficiency at antagonizing the original gene was evolutionarily refined over the mutagenesis phase which took place after the duplication event (about 2.4 mya). In addition, but independently from their insolubility, the SRGAP2A:SRGAP2C heterodimers present a significantly reduced ability to bind *ROBO1* [41].

SRGAP2A mRNA has been shown to be expressed in different regions of the central nervous system at early developmental stages. It was found to be expressed at embryonic and postnatal days in many tissues in mice, including the dorsal and ventral thalamus, the ventrolateral thalamic nucleus, the superior and inferior colliculi, the cerebellum, and the spinal cord [61]. Also in mice, Guerrier et al. [65] detected that it follows an increasing pattern of expression during early development in the cortex, reaching its maximum level at postnatal day 1 (P1), then stabilizing until P15, and gradually decreasing although still being expressed in adult stages. Charrier et al. [66] compare its expression with that of *SRGAP2C* and reach the conclusion that both are expressed in embryonic and adult human brain (though not always in exactly the same way). Various human brain expression databases we consulted^1^ generally agree that SRGAPs are expressed in frontal parts of the neocortex early in development. (Data on *SRGAP2C* specifically tend to be too sparse to draw any firm conclusion at this point.)

## 3 Concluding remarks

*SRGAP2C* may have had other functional consequences [66, 65, 68], but we have provided evidence that mechanistically we can expect *SRGAP2C* to have had an effect on the SLIT-ROBO axon guidance pathway, and — no doubt together with other genetic changes – may have contributed to the establishment of a critical aspect of the vocal learning circuit, as first suggested in [15]. We have shown that until very recently studies focusing on *SRGAP2* failed to provide evidence in this direction. It is only thanks to the results in [64, 41] and the link between filopodia and axon guidance made precise in [72] that we can adduce a greater degree of plausibility to the claim in [15] that *SRGAP2* duplications may have contributed to the emergence of aspects of our language faculty, a claim made at a time when the relevant results we rely on had not yet been obtained. Since paleogenomic work has shown that the relevant mutation that led to this effect is not specific to *Homo sapiens*, we are led to conclude that core ingredients of the vocal learning pathway pre-dated the emergence of our species.

In a certain sense, *SRGAP2C* acts like the member of the SRGAP family that most closely interacts with *ROBO1*: *SRGAP1*. Unlike *SRGAP2A*, which as we saw, induces filopodia-like membrane protrusions, *SRGAP1*’s F-BAR domain prevents filopodia [69]. By inhibiting the ability of *SRGAP2A* to induce filopodia, *SRGAP2C* makes *SRGAP2* function like *SRGAP1*. In light of this, it is noteworthy that a gene expression study [73] carried out in human developing neocortical neurons has shown a relation between *ROBO1* and *SRGAP1*. Both genes were found to be coexpressed in human corticospinal axons at various fetal periods during the formation of the corticospinal tract, which is the main descending sensorimotor projection, an elaboration of which could have given rise to the critical connection of the posterior vocal learning circuit.

As pointed out in [52], *SLIT1* is a direct target of *FOXP2* [74, 75]. Although human *FOXP2* has been reported to modulate stronger upregulation of *SLIT1* than chimpanzee *FOXP2* [75], which does not fit well with the relevant convergent downregulation of *SLIT1* in vocal learning birds found in [52], *SLIT1* is among the *FOXP2* targets found to be significantly downregulated in response to *FOXP2* expression in [76]. So, there could be another synergistic effect here between the effect of *FOXP2* on *SLIT1* and the action of *SRGAP2C* on the SLIT-ROBO pathway.

Incidentally, just like *SRGAP2C* works its effect on the SLIT-ROBO pathway by inhibiting an inhibitor (in this case, *SRGAP2A*), *FOXP2* also appears to work its effects by inhibiting inhibitors, such as *MEF2C*. As reported in [77], (mouse) *Foxp2* controls synaptic wiring of corticostriatal circuits, critical for vocal learning, by opposing *Mef2c*, which itself suppresses corticostriatal synapse formation and striatal spinogenesis. So, achieving a positive effect (establishment of a vocal learning circuit) by inhibiting inhibitors or suppressing the activity of suppressors, appear to have been a common strategy in the evolution of our lineage and our cognitive phenotype.

We still don’t know exactly when the relevant *FOXP2* mutations emerged in our lineage, so we cannot know for sure if the emergence of modern *SRGAP2C* coincided with the two *FOXP2* mutations thought to be critical for vocal learning. Evidence for a selective sweep associated with *FOXP2* yields ambiguous results (assuming that the relevant mutations were the actual selection targets): there is evidence for a recent, *H. sapiens* specific partial selective sweep [78, 79], but also evidence for another, much earlier sweep [79, Suppl. Table S12.1].

It remains to be seen if these sweeps correspond to landmarks in the establishment of the human vocal learning circuit, possibly corresponding to the stages that can be derived from the work on vocal learning birds (e.g., suboscine/proto-vocal-learning stage [37], core vocal learning circuit stage [52], shell vocal learning circuit stage [15]).

Though modest, we think that our contribution is of a kind that is necessary to make claims about when components of our language faculty mosaic emerged. It won’t do to simply identify changes on potentially relevant genes. It is necessary to show that the changes have functional effects of the right kind. We hope to have taken a small step in this direction.

## Footnotes

1 Brainspan (http://www.brainspan.org), Human Brain Transcriptome (http://hbatlas.org), Bgee (http://bgee.org), Proteomics DB (https://proteomicsdb.org), Human Protein Atlas (http://www.proteinatlas.org), Gene Enrichment Profiler (http://xavierlab2.mgh.harvard.edu/EnrichmentProfiler/index.html), and GTex (http://www.gtexportal.org).

## Author contributions statement

CB formulated the hypothesis and directed the study. PTM, MM, and CB reviewed the literature, and wrote the article. The authors declare no conflict of interest.

## References

[1] Derek Bickerton. From protolanguage to language: The speciation of modern Homo sapiens. Oxford University Press, Oxford, 2002.

[2] Steven Mithen. The singing Neanderthals: The origin of language, music, mind and body. Weidenfeld and Nicolson, London, 2005.

[3] Noam Chomsky. Some simple evo devo theses: how true might they be for language? In The Evolution of Human Language: Biolinguistic Perspectives. Cambridge University Press. 2010.

[4] Dan Dediu and Stephen C Levinson. On the antiquity of language: the reinterpretation of neandertal linguistic capacities and its consequences. Frontiers in psychology, 4:397, 2013.

[5] W Tecumseh Fitch, editor. Special Issue on the Biology and Evolution of Language (Psychonomic Bulletin & Review). Springer, 2017.

[6] M J West-Eberhard. Developmental Plasticity and Evolution. Oxford University Press, Oxford, 2003.

[7] Cedric Boeckx. Biolinguistics: forays into human cognitive biology. J. Anthropol. Sci, 91:63–89, 2013.

[8] W Tecumseh Fitch. Empirical approaches to the study of language evolution. Psychonomic Bulletin & Review, 24(1):3–33, 2017.

[9] Svante Pääbo. Neanderthal man: In search of lost genomes. Basic Books, New York, NY, 2014.

[10] Johannes Krause, Carles Lalueza-Fox, Ludovic Orlando, Wolfgang Enard, Richard E Green, Hernán A Burbano, Jean-Jacques Hublin, Catherine Hänni, Javier Fortea, Marco De La Rasilla, et al. The derived *foxp2* variant of modern humans was shared with neandertals. Current biology, 17(21):1908–1912, 2007.

[11] Cecilia SL Lai, Simon E Fisher, Jane A Hurst, Faraneh Vargha-Khadem, and Anthony P Monaco. A forkhead-domain gene is mutated in a severe speech and language disorder. Nature, 413(6855):519–523, 2001.

[12] Wolfgang Enard, Sabine Gehre, Kurt Hammerschmidt, Sabine M Hölter, Torsten Blass, Mehmet Somel, Martina K Brückner, Christiane Schreiweis, Christine Winter, Reinhard Sohr, et al. A humanized version of Foxp2 affects cortico-basal ganglia circuits in mice. Cell, 137(5):961–971, 2009.

[13] Christiane Schreiweis, Ulrich Bornschein, Eric Burguière, Cemil Kerimoglu, Sven Schreiter, Michael Dannemann, Shubhi Goyal, Ellis Rea, Catherine A French, Rathi Puliyadi, et al. Humanized Foxp2 accelerates learning by enhancing transitions from declarative to procedural performance. Proceedings of the National Academy of Sciences, 111(39):14253–14258, 2014.

[14] Fernando L Mendez, G David Poznik, Sergi Castellano, and Carlos D Bustamante. The divergence of neandertal and modern human Y chromosomes. The American Journal of Human Genetics, 98(4):728–734, 2016.

[15] Mukta Chakraborty, Solveig Walløe, Signe Nedergaard, Emma E Fridel, Torben Dabelsteen, Bente Pakkenberg, Mads F Bertelsen, Gerry M Dorrestein, Steven E Brauth, Sarah E Durand, et al. Core and shell song systems unique to the parrot brain. PLoS One, 10(6):e0118496, 2015.

[16] Dieter G Hillert. On the evolving biology of language. Frontiers in psychology, 6:1796, 2015.

[17] Christopher I Petkov and Erich Jarvis. Birds, primates, and spoken language origins: behavioral phenotypes and neurobiological substrates. Frontiers in evolutionary neuroscience, 4:12, 2012.

[18] Helen H. Shen. News feature: Singing in the brain. Proceedings of the National Academy of Sciences, 114(36):9490–9493, 2017.

[19] Erich D Jarvis and Claudio V Mello. Molecular mapping of brain areas involved in parrot vocal communication. Journal of Comparative Neurology, 419(1):1–31, 2000.

[20] Erich D Jarvis. Learned birdsong and the neurobiology of human language. Annals of the New York Academy of Sciences, 1016(1):749–777, 2004.

[21] Andreas R Pfenning, Erina Hara, Osceola Whitney, Miriam V Rivas, Rui Wang, Petra L Roulhac, Jason T Howard, Morgan Wirthlin, Peter V Lovell, Ganeshkumar Ganapathy, et al. Convergent transcriptional specializations in the brains of humans and song-learning birds. Science, 346(6215):1256846, 2014.

[22] Gesa Feenders, Miriam Liedvogel, Miriam Rivas, Manuela Zapka, Haruhito Horita, Erina Hara, Kazuhiro Wada, Henrik Mouritsen, and Erich D Jarvis. Molecular mapping of movement-associated areas in the avian brain: a motor theory for vocal learning origin. PLoS One, 3(3):e1768, 2008.

[23] Kristina Simonyan. The laryngeal motor cortex: its organization and connectivity. Current opinion in neurobiology, 28:15–21, 2014.

[24] Erich D Jarvis, Jing Yu, Miriam V Rivas, Haruhito Horita, Gesa Feenders, Osceola Whitney, Syrus C Jarvis, Electra R Jarvis, Lubica Kubikova, Ana EP Puck, et al. Global view of the functional molecular organization of the avian cerebrum: mirror images and functional columns. Journal of Comparative Neurology, 521(16):3614–3665, 2013.

[25] Yuan Wang, Agnieszka Brzozowska-Prechtl, and Harvey J Karten. Laminar and columnar auditory cortex in avian brain. Proceedings of the National Academy of Sciences, 107(28):12676–12681, 2010.

[26] Toru Shimizu, Tadd B Patton, and Scott A Husband. Avian visual behavior and the organization of the telencephalon. Brain, behavior and evolution, 75(3):204–217, 2010.

[27] W Tecumseh Fitch, Ludwig Huber, and Thomas Bugnyar. Social cognition and the evolution of language: constructing cognitive phylogenies. Neuron, 65(6):795–814, 2010.

[28] W Tecumseh Fitch. The evolution of syntax: an exaptationist perspective. Front. Evol. Neurosci, 3(9), 2011.

[29] Mukta Chakraborty and Erich D Jarvis. Brain evolution by brain pathway duplication. Phil. Trans. R. Soc. B, 370(1684):20150056, 2015.

[30] Wilder Penfield and Edwin Boldrey. Somatic motor and sensory representation in the cerebral cortex of man as studied by electrical stimulation. Brain: A journal of neurology, 1937.

[31] Ralph MW Rödel, Arno Olthoff, Frithjof Tergau, Kristina Simonyan, Dorit Kraemer, Holger Markus, and Eberhard Kruse. Human cortical motor representation of the larynx as assessed by transcranial magnetic stimulation (TMS). The Laryngoscope, 114(5):918–922, 2004.

[32] Alexander SF Leyton and Charles S Sherrington. Observations on the excitable cortex of the chimpanzee, orang-utan, and gorilla. Experimental Physiology, 11(2):135–222, 1917.

[33] MH Hast, JM Fischer, AB Wetzel, and VE Thompson. Cortical motor representation of the laryngeal muscles in macaca mulatta. Brain research, 73(2):229–240, 1974.

[34] Oscar Sugar, Joseph G Chusid, and JOHN D French. A second motor cortex in the monkey (macaca mulatta). Journal of neuropathology and experimental neurology, 7(2):182–189, 1948.

[35] Malcolm H Hast and Rosko Milojkvic. The response of the vocal folds to electrical stimulation of the inferior frontal cortex of the squirrel monkey. Acta oto-laryngologica, 61(1–6):196–204, 1966.

[36] Michel Belyk and Steven Brown. The origins of the vocal brain in humans. Neuroscience & Biobehavioral Reviews, 2017.

[37] Wan-chun Liu, Kazuhiro Wada, Erich D Jarvis, and Fernando Nottebohm. Rudimentary substrates for vocal learning in a suboscine. Nature communications, 4, 2013.

[38] Leo Joseph, Alicia Toon, Erin E Schirtzinger, Timothy F Wright, and Richard Schodde. A revised nomenclature and classification for family-group taxa of parrots (psittaciformes). Zootaxa, 3205(2), 2012.

[39] Cedric Boeckx. Language evolution. In Jon Kaas, editor, Evolution of Nervous Systems, 2nd ed., vol. 4, pages 325–339. Elsevier, London, 2017.

[40] Megan Y Dennis, Xander Nuttle, Peter H Sudmant, Francesca Antonacci, Tina A Graves, Mikhail Nefedov, Jill A Rosenfeld, Saba Sajjadian, Maika Malig, Holland Kotkiewicz, et al. Evolution of human-specific neural srgap2 genes by incomplete segmental duplication. Cell, 149(4):912–922, 2012.

[41] Michael Sporny, Julia Guez-Haddad, Annett Kreusch, Sivan Shakartzi, Avi Neznansky, Alice Cross, Michail N Isupov, Britta Qualmann, Michael M Kessels, and Yarden Opatowsky. Structural history of human SRGAP2 proteins. Molecular biology and evolution, 34.

[42] Michel Brunet, Franck Guy, David Pilbeam, Hassane Taisso Mackaye, Andossa Likius, Djimdoumalbaye Ahounta, Alain Beauvilain, Cécile Blondel, Hervé Bocherens, Jean-Renaud Bois-serie, et al. A new hominid from the Upper Miocene of Chad, Central Africa. Nature, 418(6894):145–151, 2002.

[43] Michel Brunet, Franck Guy, David Pilbeam, Daniel E Lieberman, Andossa Likius, Hassane T Mackaye, Marcia S Ponce de Leon, Christoph PE Zollikofer, and Patrick Vignaud. New material of the earliest hominid from the Upper Miocene of Chad. Nature, 434(7034):752–755, 2005.

[44] Patrick Vignaud, Philippe Duringer, Hassane Taïsso Mackaye, Andossa Likius, Cécile Blondel, Jean-Renaud Boisserie, Louis de Bonis, Véra Eisenmann, Marie-Esther Etienne, Denis Geraads, et al. Geology and palaeontology of the Upper Miocene Toros-Menalla hominid locality, Chad. Nature, 418(6894):152–155, 2002.

[45] Nick Patterson, Daniel J Richter, Sante Gnerre, Eric S Lander, and David Reich. Genetic evidence for complex speciation of humans and chimpanzees. Nature, 441(7097):1103–1108, 2006.

[46] Dieter Hillert. The premodern language-ready brain of our common ancestor. Ms. University of California, San Diego, 2015.

[47] Randy L Buckner and Fenna M Krienen. The evolution of distributed association networks in the human brain. Trends in Cognitive Sciences, 17(12):648–665, 2013.

[48] Suzana Herculano-Houzel. The remarkable, yet not extraordinary, human brain as a scaled-up primate brain and its associated cost. Proceedings of the National Academy of Sciences, 109(Supplement 1):10661–10668, 2012.

[49] Suzana Herculano-Houzel. The Human Advantage. MIT Press, Cambridge, MA, 2016.

[50] Basant K Tiwary. Evolution of the SRGAP2 gene is linked to intelligence in mammals. Biomedicine Hub, 1(1):443947–443947, 2016.

[51] Barry J Dickson. Molecular mechanisms of axon guidance. Science, 298(5600):1959–1964, 2002.

[52] Rui Wang, Chun-Chun Chen, Erina Hara, Miriam V Rivas, Petra L Roulhac, Jason T Howard, Mukta Chakraborty, Jean-Nicolas Audet, and Erich D Jarvis. Convergent differential regulation of SLIT-ROBO axon guidance genes in the brains of vocal learners. Journal of Comparative Neurology, 523(6):892–906, 2015.

[53] Katja Brose, Kimberly S Bland, Kuan Hong Wang, David Arnott, William Henzel, Corey S Goodman, Marc Tessier-Lavigne, and Thomas Kidd. Slit proteins bind Robo receptors and have an evolutionarily conserved role in repulsive axon guidance. Cell, 96(6):795–806, 1999.

[54] Hua Long, Christelle Sabatier, Le Ma, Andrew Plump, Wenlin Yuan, David M Ornitz, Atsushi Tamada, Fujio Murakami, Corey S Goodman, and Marc Tessier-Lavigne. Conserved roles for Slit and Robo proteins in midline commissural axon guidance. Neuron, 42(2):213–223, 2004.

[55] Pontus Aspenström. Roles of f-bar/pch proteins in the regulation of membrane dynamics and actin reorganization. International review of cell and molecular biology, 272:1–31, 2008.

[56] Kit Wong, Xiu-Rong Ren, Yang-Zhong Huang, Yi Xie, Guofa Liu, Harumi Saito, Hao Tang, Leng Wen, Susann M Brady-Kalnay, Lin Mei, et al. Signal transduction in neuronal migration: roles of GTPase activating proteins and the small GTPase Cdc42 in the Slit-Robo pathway. Cell, 107(2):209–221, 2001.

[57] Alan Hall. Rho GTPases and the actin cytoskeleton. Science, 279(5350):509–514, 1998.

[58] Qin Yao, Wei-Lin Jin, Ying Wang, and Gong Ju. Regulated shuttling of Slit-Robo-GTPase activating proteins between nucleus and cytoplasm during brain development. Cellular and molecular neurobiology, 28(2):205–221, 2008.

[59] Valérie Marillat, Oliver Cases, Kim Tuyen Nguyenf-Ba-Charvet, Marc Tessier-Lavigne, Constantino Sotelo, and Alain Chédotal. Spatiotemporal expression patterns of slit and robo genes in the rat brain. Journal of Comparative Neurology, 442(2):130–155, 2002.

[60] Kristin L Whitford, Valérie Marillat, Elke Stein, Corey S Goodman, Marc Tessier-Lavigne, Alain Chédotal, and Anirvan Ghosh. Regulation of cortical dendrite development by Slit-Robo interactions. Neuron, 33(1):47–61, 2002.

[61] Claire Bacon, Volker Endris, and Gudrun Rappold. Dynamic expression of the Slit-Robo GTPase activating protein genes during development of the murine nervous system. Journal of Comparative Neurology, 513(2):224–236, 2009.

[62] Xiaofeng Li, Yushu Chen, Yiwei Liu, Jia Gao, Feng Gao, Mark Bartlam, Jane Y Wu, and Zihe Rao. Structural basis of Robo proline-rich motif recognition by the srGAP1 Src homology 3 domain in the Slit-Robo signaling pathway. Journal of Biological Chemistry, 281(38):28430–28437, 2006.

[63] Hirokazu Okada, Akiyoshi Uezu, Frank M Mason, Erik J Soderblom, M Arthur Moseley III, and Scott H Soderling. SH3 Domain–Based Phototrapping in Living Cells Reveals Rho Family GAP Signaling Complexes. Science signaling, 4(201):rs13, 2011.

[64] Julia Guez-Haddad, Michael Sporny, Yehezkel Sasson, Lada Gevorkyan-Airapetov, Naama Lahav-Mankovski, David Margulies, Jens Radzimanowski, and Yarden Opatowsky. The neu-ronal migration factor srGAP2 achieves specificity in ligand binding through a two-component molecular mechanism. Structure, 23(11):1989–2000, 2015.

[65] Sabrice Guerrier, Jaeda Coutinho-Budd, Takayuki Sassa, Aurélie Gresset, Nicole Vincent Jordan, Keng Chen, Wei-Lin Jin, Adam Frost, and Franck Polleux. The F-BAR domain of srGAP2 induces membrane protrusions required for neuronal migration and morphogenesis. Cell, 138(5):990–1004, 2009.

[66] Cécile Charrier, Kaumudi Joshi, Jaeda Coutinho-Budd, Ji-Eun Kim, Nelle Lambert, Jacqueline De Marchena, Wei-Lin Jin, Pierre Vanderhaeghen, Anirvan Ghosh, Takayuki Sassa, et al. Inhibition of SRGAP2 function by its human-specific paralogs induces neoteny during spine maturation. Cell, 149(4):923–935, 2012.

[67] Ruth Benavides-Piccione, Inmaculada Ballesteros-Yáñez, Javier DeFelipe, and Rafael Yuste. Cortical area and species differences in dendritic spine morphology. Journal of neurocytology, 31(3):337–346, 2002.

[68] Matteo Fossati, Rocco Pizzarelli, Ewoud R Schmidt, Justine V Kupferman, David Stroebel, Franck Polleux, and Cécile Charrier. SRGAP2 and its human-specific paralog co-regulate the development of excitatory and inhibitory synapses. Neuron, 91(2):356–369, 2016.

[69] Jaeda Coutinho-Budd, Vladimir Ghukasyan, Mark J Zylka, and Franck Polleux. The F-BAR domains from srGAP1, srGAP2 and srGAP3 regulate membrane deformation differently. J Cell Sci, 125(14):3390–3401, 2012.

[70] Rafael Dominik Fritz, Denis Menshykau, Katrin Martin, Andreas Reimann, Valeria Pontelli, and Olivier Pertz. SrGAP2-dependent integration of membrane geometry and Slit-Robo-repulsive cues regulates fibroblast contact inhibition of locomotion. Developmental cell, 35(1):78–92, 2015.

[71] Erik W Dent and Frank B Gertler. Cytoskeletal dynamics and transport in growth cone motility and axon guidance. Neuron, 40(2):209–227, 2003.

[72] Russell E McConnell, J Edward van Veen, Marina Vidaki, Adam V Kwiatkowski, Aaron S Meyer, and Frank B Gertler. A requirement for filopodia extension toward Slit during Robo-mediated axon repulsion. J Cell Biol, 213(2):261–274, 2016.

[73] Bui Kar Ip, Nadhim Bayatti, Nicholas J Howard, Susan Lindsay, and Gavin J Clowry. The corticofugal neuron-associated genes ROBO1, SRGAP1, and CTIP2 exhibit an anterior to posterior gradient of expression in early fetal human neocortex development. Cerebral Cortex, 21(6):1395–1407, 2011.

[74] Sonja C Vernes, Elizabeth Spiteri, Jérôme Nicod, Matthias Groszer, Jennifer M Taylor, Kay E Davies, Daniel H Geschwind, and Simon E Fisher. High-throughput analysis of promoter occupancy reveals direct neural targets of FOXP2, a gene mutated in speech and language disorders. The American Journal of Human Genetics, 81(6):1232–1250, 2007.

[75] Genevieve Konopka, Jamee M Bomar, Kellen Winden, Giovanni Coppola, Zophonias O Jonsson, Fuying Gao, Sophia Peng, Todd M Preuss, James A Wohlschlegel, and Daniel H Geschwind. Human-specific transcriptional regulation of CNS development genes by FOXP2. Nature, 462(7270):213–217, 2009.

[76] Paolo Devanna, Jeroen Middelbeek, and Sonja C Vernes. FOXP2 drives neuronal differentiation by interacting with retinoic acid signaling pathways. Frontiers in cellular neuroscience, 8:305, 2014.

[77] Yi-Chuan Chen, Hsiao-Ying Kuo, Ulrich Bornschein, Hiroshi Takahashi, Shih-Yun Chen, Kuan-Ming Lu, Hao-Yu Yang, Gui-May Chen, Jing-Ruei Lin, Yi-Hsin Lee, et al. Foxp2 controls synaptic wiring of corticostriatal circuits and vocal communication by opposing Mef2c. Nature Neuroscience, 2016.

[78] Tomislav Maricic, Viola Günther, Oleg Georgiev, Sabine Gehre, Marija Ćurlin, Christiane Schreiweis, Ronald Naumann, Hernán A Burbano, Matthias Meyer, Carles Lalueza-Fox, et al. A recent evolutionary change affects a regulatory element in the human FOXP2 gene. Molecular Biology and Evolution, 30(4):844–852, 2013.

[79] Swapan Mallick, Heng Li, Mark Lipson, Iain Mathieson, Melissa Gymrek, Fernando Racimo, Mengyao Zhao, Niru Chennagiri, Susanne Nordenfelt, Arti Tandon, et al. The simons genome diversity project: 300 genomes from 142 diverse populations. Nature, 538(7624):201–206, 2016.

